# The DUSP1 on CaMKII-positive neurons in mPFC mediates adolescent cocaine exposure-induced higher sensitivity to cocaine during adulthood

**DOI:** 10.1101/2022.08.26.505502

**Authors:** Xiaoyan Wei, Jiasong Chang, Zhen Cheng, Wenwen Chen, Hao Guo, Zhaoyu Liu, Yuning Mai, Tao Hu, Yuanyuan Zhang, Qinglong Cai, Feifei Ge, Yu Fan, Xiaowei Guan

**Affiliations:** Department of Human Anatomy and Histoembryology, Nanjing University of Chinese Medicine, Nanjing, China

**Keywords:** adolescent cocaine exposure, sensitivity to cocaine, mPFC activation, DUSP1^CaMKII^

## Abstract

Adolescent cocaine abuse increases the risk for developing addiction in later life, which remains a big public health concern, but the underlying molecular mechanism is poorly understood. Here, adolescent cocaine-exposed (ACE) male mice models were established by administrating cocaine during adolescent period. When growing to adult age, mice were subjected to conditioned place preference (CPP) to evaluate the sensitivity to cocaine, then potential molecule of dual specificity phosphatase 1 (DUSP1) were screened out by transcriptomic sequencing. Subthreshold dose of cocaine (sdC), that is insufficient to produce CPP, was used to induce CPP in adulthood. The sdC treatment effectively induced CPP in ACE mice during adulthood, accompanied with the more triggered CaMKII-positive neurons, and induced higher *Dusp1* gene, lower DUSP1 protein, lower DUSP1 activity and lower DUSP1 expression on CaMKII-positive neurons (DUSP1^CaMKII^) in medial prefrontal cortex (mPFC). Overexpressing DUSP1^CaMKII^ suppressed CaMKII-positive neuronal activation, and ultimately blocked sdC-induced CPP in ACE mice during adulthood. While, knocking-down DUSP1^CaMKII^ activated more CaMKII-positive neurons, and aggravated sdC-preferred behavior in ACE mice during adulthood. ERK1/2 might be potential subsequent signal for DUSP1 in the process. Collectively, our findings reveal a novel molecular mechanism underlying adolescent drug abuse-induced susceptibility to addiction during adulthood, and mPFC DUSP1^CaMKII^ is a promising pharmacological target to predict or treat addiction, especially caused by adolescent substance use.

**Summary:** Adolescent cocaine exposure causes higher cocaine-preferred behaviors during adulthood, along with evoked mPFC activity in response to cocaine challenge. Locally overexpressing but not knocking-down the dual specificity phosphatase 1 (DUSP1) on CaMKII-positive neurons (DUSP1^CaMKII^) suppresses mPFC activation, and ultimately rescues the higher sensitivity to cocaine during adulthood.

## Introduction

Adolescents present vulnerable to substance abuse such as cocaine (1, 2). Adolescent experiences of drug abuse enhances the risk for developing addiction later in life (3, 4), however, the underlying molecular mechanism remains much to be clarified. The medial prefrontal cortex (mPFC), a critical brain area involved in the susceptibility to drug addiction (5, 6), undergoes rapid development during adolescent period (7, 8). A single cocaine treatment alters mPFC dendritic spines in adolescent rather than adult animals (9, 10), indicating a sensitive response of adolescent mPFC to drugs (11, 12). Previously, we found that adolescent cocaine exposure (ACE) enhanced the GABAergic transmission on pyramidal neurons of mPFC (13), and produced a prolonged modification on mPFC synapses in adult rodents (14).

The dual specificity protein kinase-phosphatase-1 (DUSP1), also known as mitogen-activated protein kinase-phosphatase-1 (MKP-1), is a negative regulatory factor on the activation of mitogen activated protein kinases (MAPKs) (15). DUSP1 are expressed widely in the brain, including PFC (16). By regulating dephosphorization of MAPKs such as ERK1/2, p38 and the c-Jun N-terminal kinases (JNKs), DUSP1 has been proved to achieve neuroprotection in hearing loss (17), Huntington’s disease (18), cerebral ischemia reperfusion injury (19) and neuropathic pain (20) through its anti-inflammatory, antioxidant, mitochondrial repair and supporting axon survival functions. In addition, *Dusp1* gene are known to be activity-dependent responded to stimuli signals in the brain, thus often acting as an immediate early gene (IEGs) like *c-fos*. For example, psilocybin strongly upregulates *Dusp1* levels in PFC (21), while synthetic cathinone (substitutes for illicit psychostimulants including cocaine) produces increased *Dusp1* genes in striatum (22).

In the present study, ACE male mice model was established by exposing adolescent male mice to cocaine from postnatal day 28 (P28) to P42, a period that represents adolescent in rodents (23). When growing up to adult age, mice were subjected to conditioned place preference (CPP) to evaluate the sensitivity to drug. Subthreshold dose of cocaine, which is insufficient to produce CPP, was used to induce CPP (sdC). With method of transcriptomic sequencing and locally regulating DUSP1 on CaMKII-positive neurons (DUSP1^CaMKII^), the role of mPFC DUSP1^CaMKII^ in the sensitivity to cocaine were explored in ACE mice during adulthood.

## Results

### The subthreshold dose of cocaine produce CPP and trigger more mPFC CaMKII-positive neurons in ACE mice during adulthood

To examine the sensitivity to cocaine, mice were subjected to cocaine-induced CPP training and tests in mice during adulthood (Figure 1A). Subthreshold dose (1 mg / kg) of cocaine (sdC), which is insufficient to induce CPP in naïve mice, were used in this study. As shown in Figure 1B-D, the sdC treatment did not produce CPP in adolescent saline-exposed (ASE-sdC) mice (n = 18 mice, *t* = 1.306, *p* = 0.3586), but produced CPP in ACE mice during adulthood (ACE-sdC) (n = 24 mice, *t* = 10.91, *p* < 0.0001, Figure 1B, 1D). The ΔCPP score was higher in ACE-sdC mice than that in ASE-sdC mice (n = 42 mice, *t* = 8.129, *p* < 0.0001, Figure 1C, 1D). These results indicate that adolescent cocaine exposure produce a higher sensitivity to cocaine in mice during adulthood.

**Figure 1.**
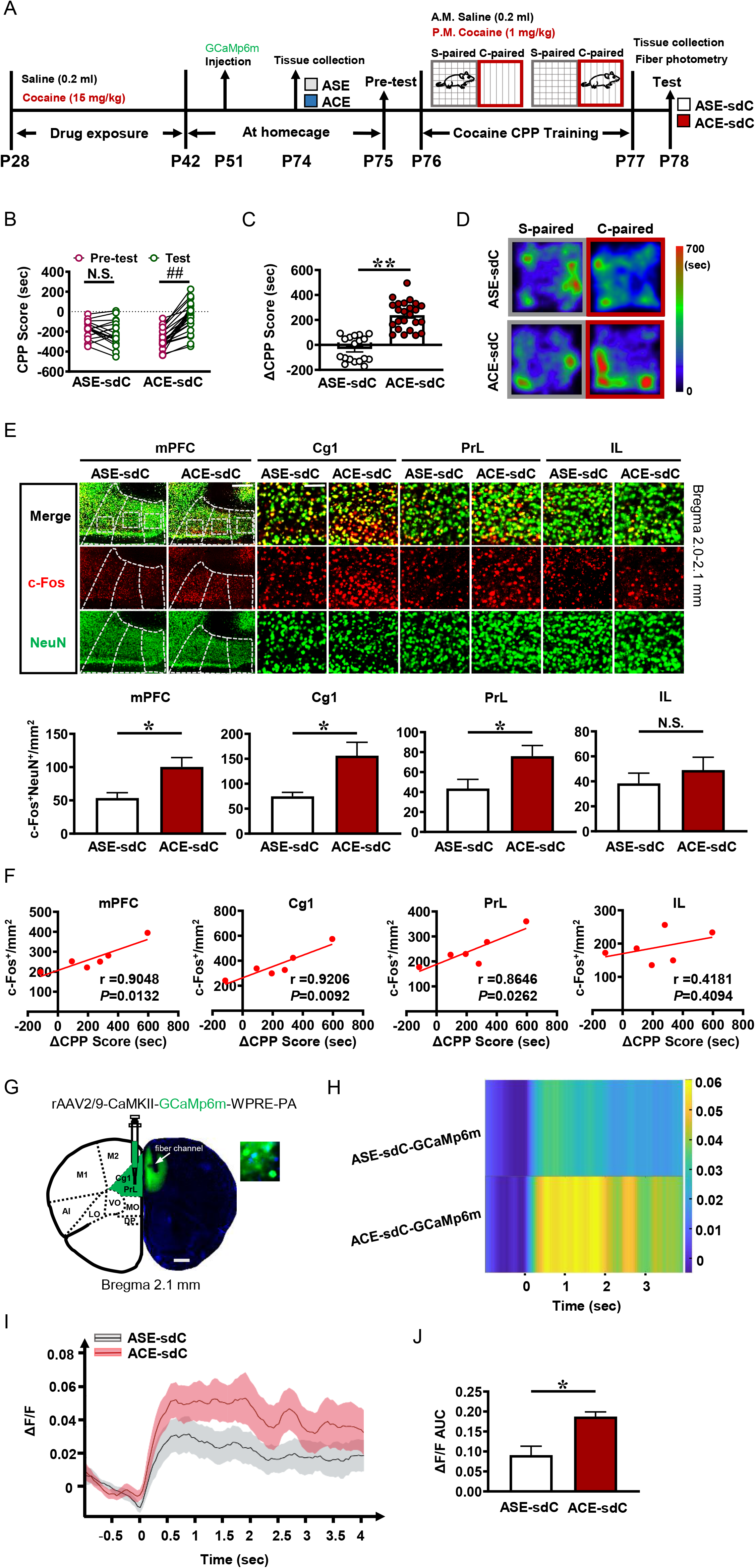
Subthreshold dose of cocaine elicit cocaine CPP and trigger mPFC CaMKII-positive neurons in ACE mice during adulthood. **A,** Experimental models and timeline. **B,** CPP score. **C,** ΔCPP score. **D,** Heatmap of spent duration by mice in CPP apparatus. **E,** C-Fos staining after CPP test in mPFC and its subregions of Cg1, PrL, IL. The c-Fos (red) and NeuN (green) are used as markers for evaluating cell activity and labeling neuron, respectively. Scale bar, 400 μm/100 μm. **F,** Schematic diagram of viral injection. **G,** CPP score. **H,** ΔCPP score. **I,** Heatmap of spent duration by mice in CPP apparatus. **J,** Heatmap of GCaMp6m fluorescence at first 4 sec when mice entering cocaine-paired chamber. **K,** Quantification of ΔF/F. **L,** The AUC of ΔF/F. Cg1, cingulate cortex; PrL, prelimbic cortex; IL, infralimbic cortex; ASE-sdC, adolescent saline-exposed mice with subthreshold dose of cocaine treatment during adulthood; ACE-sdC, adolescent cocaine-exposed mice with subthreshold dose of cocaine treatment during adulthood; N.S., *p* > 0.05, #, *p* < 0.05, ##, *p* < 0.01 vs baseline (pre-test CPP score); N.S., *p* > 0.05, *, *p* < 0.05, **, *p* < 0.01 vs ASE-sdC mice.

To map the brain activity in respond to subthreshold dose administration of cocaine in adult mice, c-Fos and NeuN were used as markers to evaluate neuronal activation and to label neuron, respectively. Compared with ASE-sdC mice during adulthood, the c-Fos-stained levels of neurons is much higher in mPFC (n = 12 slice from 6 mice, *t* = 2.89, *p* = 0.0161) and its subregions of cingulate cortex (Cg1, n = 12 slice from 6 mice, *t* = 2.912, *p* = 0.0155) and prelimbic cortex (PrL, n = 12 slice from 6 mice, *t* = 2.311, *p* = 0.0435), but similar in infralimbic cortex (IL, n = 12 slice from 6 mice, *t* = 0.8092, *p* = 0.4372, Figure 1E), orbitofrontal cortex (OFC, n = 12 slice from 6 mice, *t* = 2.055, *p* = 0.0670), nucleus accumbens core (NAc core, n = 12 slice from 6 mice, *t* = 0.3219, *p* = 0.7542), nucleus accumbens shell (NAc shell, n = 12 slice from 6 mice, *t* = 1.632, *p* = 0.1337), bed nucleus of the stria terminalis (BNST, n = 12 slice from 6 mice, *t* = 1.172, *p* = 0.2685), triangular septal nucleus (TS, n = 12 slice from 6 mice, *t* = 1.768, *p* = 0.1074), basolateral amygdala (BLA, n = 12 slice from 6 mice, *t* = 1.304, *p* = 0.2215), central nucleus of the amygdala (CeA, n = 12 slice from 6 mice, *t* = 0.2851, *p* = 0.7814), medial habenula (mHb, n = 12 slice from 6 mice, *t* = 0.2165, *p* = 0.8329), lateral habenula (LHb, n = 12 slice from 6 mice, *t* = 1.151, *p* = 0.2764), dorsal hippocampal CA1 (dCA1, n = 12 slice from 6 mice, *t* = 0.09203, *p* = 0.9285), dorsal hippocampal CA3 (dCA3, n = 12 slice from 6 mice, *t* = 0.6802, *p* = 0.5118), dorsal dentate gyrus (dDG, n = 12 slice from 6 mice, *t* = 0.5304, *p* = 0.6074), ventral hippocampal CA1 (vCA1, n = 10 slice from 6 mice, *t* = 0.1572, *p* = 0.8790) and ventral tegmental area (VTA, n = 12 slice from 6 mice, *t* = 0.2915, *p* = 0.7767, Figure S1A) in ACE-sdC mice during adulthood. Further, In ACE-sdC mice, the c-Fos staining in mPFC (n = 6 mice, r = 0.9048, *p* = 0.0132), Cg1 (n = 6 mice, r = 0.9206, *p* = 0.0092) and PrL (n = 6 mice, r = 0.8646, *p* = 0.0262), but not in IL (n = 6 mice, r = 0.4181, *p* = 0.4094) showed significant and positive correlation with the corresponding ΔCPP score in ACE-sdC mice during adulthood (Figure 1F). These results imply that mPFC activation, especially Cg1 and PrL, might be involved in the adolescent cocaine exposure-induced higher sensitivity to cocaine in mice during adulthood.

To observe the effects of adolescent cocaine exposure on baseline levels of mPFC activity, the c-Fos levels without administration of sdC were test in mice on P74. As shown in Figure S2A, before the sdC administration, the c-Fos protein levels (n = 6 mice, *t* = 5.899, *p* = 0.0041) of mPFC was higher in ACE than that in ASE mice on P74. These results indicate that adolescent cocaine exposure evoke mPFC activation in mice during adulthood.

To explore the activity of mPFC different cell types in response to sdC treatment, CaMKII, GAD67, PV and SOM were used as markers to label glutamatergic, GABAergic, PV-positive GABAergic and SOM-positive GABAergic neuron, respectively. As shown in Figure S2B, the c-Fos-stained CaMKII-positive glutamatergic neurons (n = 12 slice from 6 mice, *t* = 3.656, *p* = 0.0044), but not GAD67-positive GABAergic neurons (n = 12 slice from 6 mice, *t* = 1.180, *p* = 0.2652) and its subtypes of PV-positive (n = 12 slice from 6 mice, *t* = 0.5668, *p* = 0.5833) and SOM-positive neurons (n = 12 slice from 6 mice, *t* = 0.6119, *p* = 0.5543), were much more in numbers at mPFC of ACE-sdC mice than that in ASE-sdC mice during adulthood.

To *in vivo* record real-time calcium signals of mPFC CaMKII-positive neurons, *rAAV2/9-CaMKII-GCaMp6m-WPRE-PA* virus were injected into mPFC, specially limiting to subregions of Cg1 and PrL (Figure 1G). As shown in Figure 1H-J, at first 4 sec-duration after mice entering into the cocaine-paired chamber, the calcium signals of CaMKII-positive neurons were significantly at higher levels in GCaMp6m-transfected ACE-sdC mice than that in ASE-sdC mice (n = 6 mice, *t* = 3.839, *p* = 0.0185). These results indicate that mPFC CaMKII-positive neurons might be involved in adolescent cocaine exposure-induced higher sensitivity to cocaine in mice during adulthood.

### Subthreshold dose of cocaine increase mPFC *Dusp1* gene while decrease DUPS1 protein and activity in ACE mice during adulthood

To screen potential molecules that contribute to the higher mPFC activity in response to sdC administration, the RNA sequencing was performed in mPFC on P78 following CPP test. As shown in Figure S3A, following administration of sdC, 180 genes were upregulated and 248 genes were downregulated in mPFC of ACE-sdC mice, when compared to ASE-sdC mice. According to Protein-Protein Interaction (PPI) network analysis of upregulated genes (Figure S3B) and heatmaps of top 40 upregulated genes (Figure S3C), *Dusp1* is one of the most significantly upregulated genes and a hub interactor gene that associated with other upregulated genes including *c-Fos*. Further, MAPK-related signals were both in the top 30 enriched GO (Figure S3D) and the top enriched 20 KEGG pathway (Figure S3E) of upregulated genes in mPFC of ACE-sdC mice. The transcriptomic results imply that *Dusp1*/DSUP1 and related MAPK signals might be a critical molecule that implicated in adolescent cocaine exposure-increased sensitivity to cocaine.

The baseline level and responsive level to cocaine of mPFC *Dusp1* gene, DUSP1 protein and activity were examined on P74 and P78, respectively. There is similar *Dusp1* gene (n = 6 mice, *t* = 0.3691, *p* = 0.7307, Figure 2A), DUSP1 protein (n = 6 mice, *t* = 1.905, *p* = 0.1295, Figure 2B), and DUSP1 activity (n = 6 mice, *t* = 0.1855, *p* = 0.8618, Figure 2C) between ACE and ASE mice on P74, indicating that adolescent cocaine exposure did not influence the baseline levels of *Dusp1/DUSP1* in mPFC. However, on P78 following administration of sdC, *Dusp1* gene (n = 11 mice, *t* = 4.100, *p* = 0.0027, Figure 2D) were much higher, while DUSP1 protein level (n = 6 mice, *t* = 4.929, *p* = 0.0079, Figure 2E), DUSP1 activity (n = 12 mice, *t* = 3.240, *p* = 0.0089, Figure 2F) and DUSP1-staining value on CaMKII-positive neurons (n = 6 mice, *t* = 5.751, *p* = 0.0045, Figure 2G) were much lower in ACE-sdC mice than that in ASE-sdC mice. With these results, we further infer that *Dusp1/DUSP1* on CaMKII-positive neurons (DUSP1^CaMKII^) may contribute to the sdC-evoked mPFC CaMKII-positive neurons and higher sensitivity to cocaine in ACE mice during adulthood.

**Figure 2.**
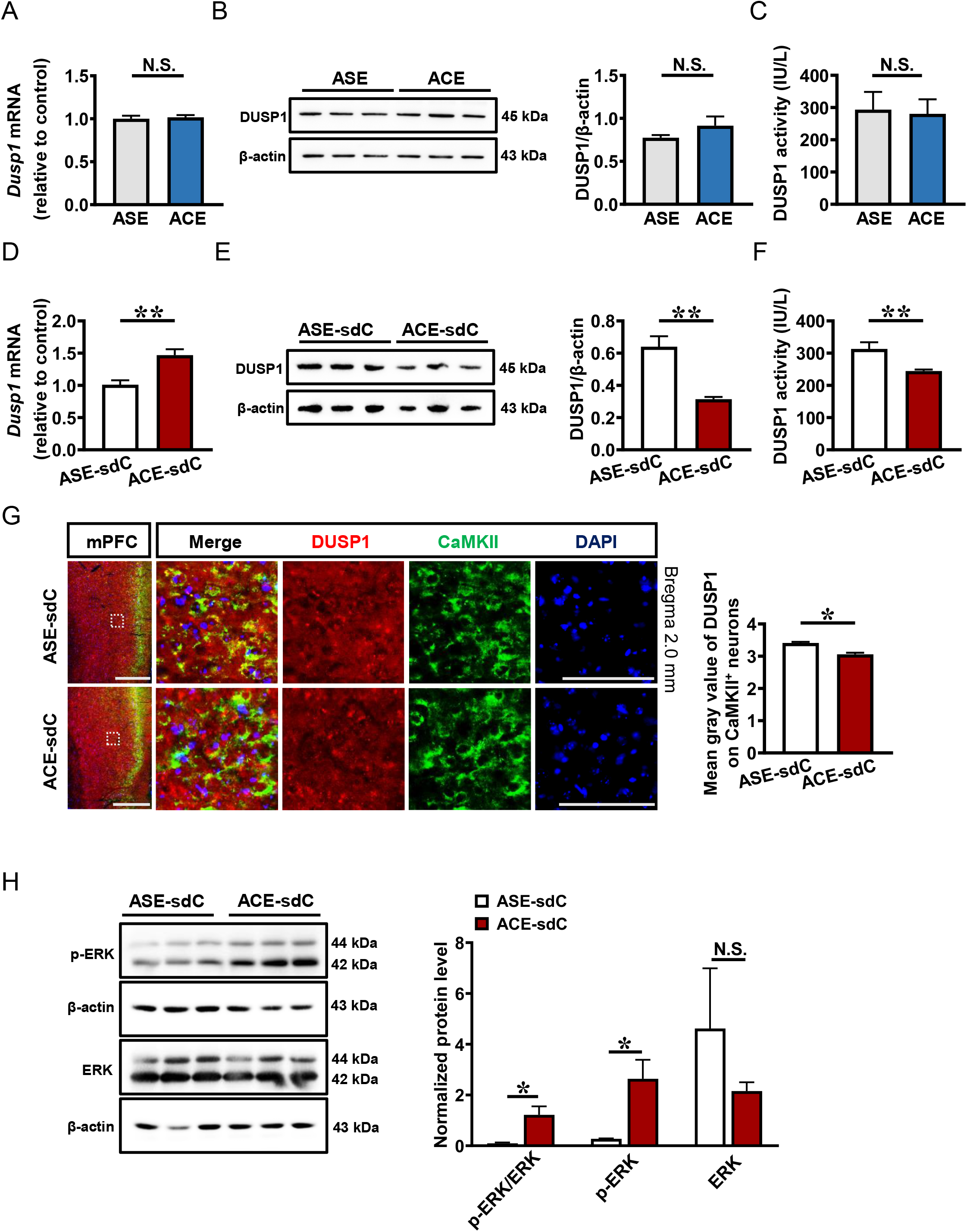
Subthreshold dose administration of cocaine increase *Dusp1* gene while decrease DUPS1 protein and activity in mPFC of adult ACE mice. **A,** *Dusp1* mRNA levels on P74. **B,** DUSP1 protein levels and activity on P74. **C,** DUSP1 activity on P74. **D,** *Dusp1* mRNA levels on P78. **E,** DUSP1 protein levels and activity on P78. **F,** DUSP1 activity on P78. **G,** DUSP1 staining in CaMKII-positive neurons. Scale bar, 400 μm/100 μm. **H,** Levels of p-ERK/ERK, p-ERK and ERK proteins on P78. ASE, adolescent saline-exposed mice during adulthood; ACE, adolescent cocaine-exposed mice during adulthood; ASE-sdC, adolescent saline-exposed mice with subthreshold dose of cocaine treatment during adulthood; ACE-sdC, adolescent cocaine-exposed mice with subthreshold dose of cocaine treatment during adulthood; N.S., *p* > 0.05, *, *p* < 0.05, **, *p* < 0.01 vs ASE mice or ASE-sdC mice.

Since DUSP1 is important regulatory molecule in the MAPK-related signal molecules, the total ERK1/2 and its phosphorylation (p-ERK) levels of mPFC was examined on P78 following sdC administration. As shown in Figure 2H, the levels of ratio of p-ERK to ERK (n = 6 mice, *t* = 3.338, *p* = 0.0289) and p-ERK (n = 6 mice, *t* = 3.143, *p* = 0.0348) were significantly higher, while total ERK1/2 (n = 6 mice, *t* = 1.030, *p* = 0.3611) was similar in ACE-sdC than that in ASE-sdC mice.

### Overexpressing DUSP1^CaMKII^ suppress cocaine-induced CPP and reduce mPFC activation in ACE mice during adulthood

To explore the role of mPFC DUSP1^CaMKII^ in the higher sensitivity to cocaine in ACE mice, *AAV9-CaMKII-DUSP1-RNAOE-EGFP* (OE virus) and *AAV9-CaMKII-EGFP* (Ctrl virus) were bilaterally infused into mPFC, especially to Cg1 and PrL (Figure 3A-B). As shown in Figure 3C, about 80% CaMKII-positive neurons were transfected with virus. Compared to Ctrl virus-treated ACE-sdC mice (ACE-sdC_Ctrl_), *Dusp1* gene (n = 6 mice, *t* = 4.413, *p* = 0.0116, Figure 3D), DUSP1 protein (n = 6 mice, *t* = 4.071, *p* = 0.0152, Figure 3E) and DUSP1 activity (n = 12 from 6 mice, *t* = 3.336, *p* = 0.0075, Figure 3F) levels were much higher in OE virus-treated ACE-sdC mice (ACE-sdCOE), indicating the good efficiency of OE virus.

**Figure 3.**
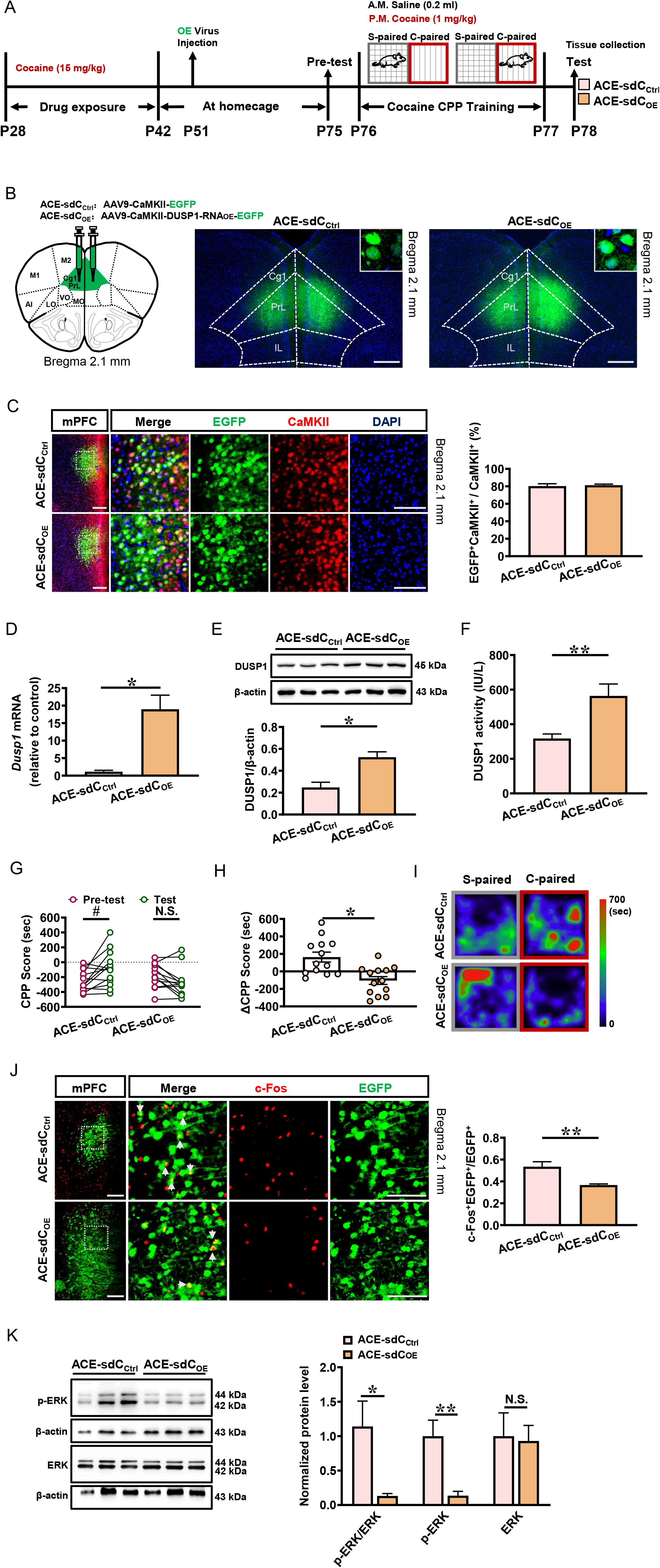
Overexpression of DUSP1^CaMKII^ reduce cocaine-induced CPP and attenuate mPFC activation in ACE mice during adulthood. **A,** Experimental design and timeline. **B,** Schematic diagram of viral injection. Scale bar, 400 μm. **C,** Percentage of viral transfected neurons in CaMKII-positive neurons of mPFC. Scale bar, 200 μm/100 μm. **D,** *Dusp1* mRNA levels on P78. **E,** DUSP1 protein levels on P78. **F,** DUSP1 activity on P78. **G,** CPP score. **H,** ΔCPP score. **I,** Heatmap of spent duration by mice in CPP apparatus. **J,** The percentage of c-Fos-positive neurons in virus-transfected neurons. Scale bar, 200 μm/100 μm. **K,** Levels of p-ERK/ERK, p-ERK and ERK proteins on P78. ACE-sdC_Ctrl_, ACE-sdC mice injected with Ctrl virus; ACE-sdCOE, ACE-sdC mice with overexpression virus of DUSP1; N.S., *p* > 0.05, #, *p* < 0.05, ##, *p* < 0.01 vs baseline (pre-test CPP score); N.S., *p* > 0.05, *, *p* < 0.05, **, *p* < 0.01 vs ACE-sdC_Ctrl_ mice.

As shown in Figure 3G-I, ACE-sdC_Ctrl_ mice still showed sdC-induced preference behavior (n = 13 mice, *t* = 3.215, *p* = 0.0074), while ACE-sdCOE mice did not exhibit sdC-related CPP (n = 13 mice, *t* = 2.051, *p* = 0.1001), accompanied by a lower ΔCPP score in ACE-sdCOE mice (n = 26 mice, *t* = 3.723, *p* = 0.0011). These results indicate overexpressing DUSP1^CaMKII^ could suppress adolescent cocaine exposure-induced higher sensitivity to cocaine during adulthood.

To explore the regulatory role of DUSP1 in the activity of transfected CaMKII neurons (EGFP-positive), c-Fos staining were performed on mPFC. As shown in Figure 3J, the percentage of c-Fos-positive & EGFP-positive in transfected CaMKII neurons was much lower in ACE-sdCOE mice than that in ACE-sdCCtrl mice (n = 12 slice from 6 mice, *t* = 3.624, *p* = 0.0047). In parallel, the ratio of p-ERK to ERK (n = 6 mice, *t* = 3.503, *p* = 0.0248) and p-ERK (n = 6 mice, *t* = 9.063, *p* = 0.0008) levels were significantly decreased, but ERK level (n = 6 mice, *t* = 0.1728, *p* = 0.8712) was not changed by OE virus, compared to Ctrl virus (Figure 3K). These results indicate DUSP1^CaMKII^ negatively regulate mPFC CaMKII-positive neurons.

### Knocking-down DUSP1^CaMKII^ aggravate cocaine-preferred behaviors and further activate mPFC activity in ACE mice during adulthood

To explore the role of ACE-increased *Dusp1* gene in further confirm the role of mPFC DUSP1^CaMKII^ in mPFC activity and sensitivity to cocaine, *AAV9-CaMKII-DUSP1-RNAi-EGFP* (KD virus) and *AAV9-CaMKII-EGFP* (Ctrl virus) were bilaterally infused into mPFC, especially to Cg1 and PrL (Figure 4A-B). As shown in Figure 4C, above 70% CaMKII-positive neurons were transfected with virus. Compared to Ctrl virus-treated ACE-sdC mice (ACE-sdCCtrl mice), *Dusp1* gene (n = 8 mice, *t* = 9.250, *p* < 0.0001, Figure 4D), DUSP1 protein (n = 6 mice, *t* = 11.00, *p* = 0.0004, Figure 4E) and DUSP1 activity (n = 11 from 6 mice, *t* = 3.256, *p* = 0.0099, Figure 4F) were lower in KD virus-treated ACE-sdC mice (ACE-sdC_KD_ mice), indicating good efficiency of KD virus.

**Figure 4.**
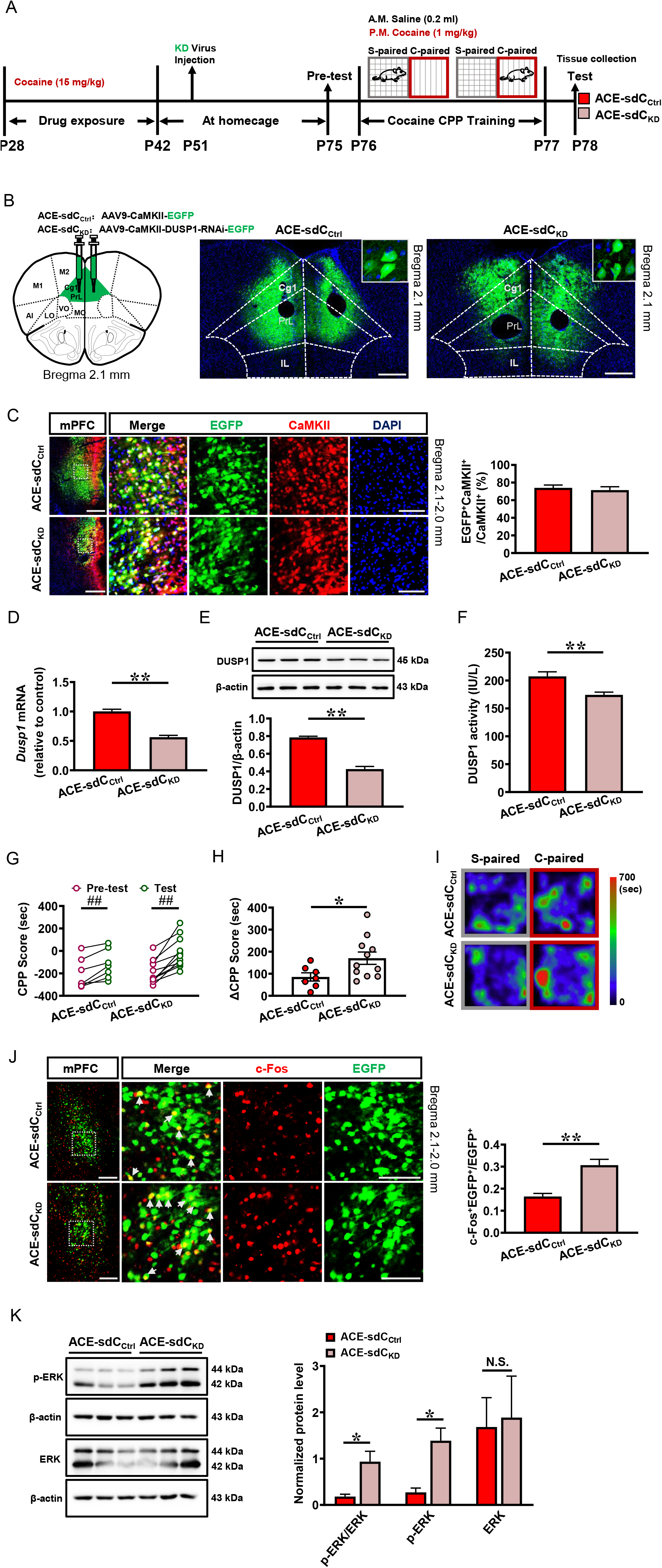
Knockdown of DUSP1^CaMKII^ aggravate cocaine-induced CPP and further activate mPFC activity in ACE mice during adulthood. **A,** Experimental design and timeline. **B,** Schematic diagram of viral injection. Scale bar, 400 μm. **C,** Percentage of viral transfected neurons in CaMKII-positive neurons of mPFC. Scale bar, 200 μm/100 μm. **D,** *Dusp1* mRNA levels on P78. **E,** DUSP1 protein levels on P78. **F,** DUSP1 activity on P78. **G,** CPP score. **H,** ΔCPP score. **I,** Heatmap of spent duration by mice in CPP apparatus. **J,** The percentage of c-Fos-positive neurons in virus-transfected neurons. Scale bar, 200 μm/100 μm. **K,** Levels of p-ERK/ERK, p-ERK and ERK proteins on P78. ACE-sdCCtrl, ACE-sdC mice injected with Ctrl virus; ACE-sdCKD, ACE-sdC mice with knockdown virus of DUSP1; N.S., *p* > 0.05, #, *p* < 0.05, ##, *p* < 0.01 vs baseline (pre-test CPP score); N.S., *p* > 0.05, *, *p* < 0.05, **, *p* < 0.01 vs ACE-sdC_Ctrl_ mice.

As shown in Figure 4G-I, both ACE-sdCCtrl mice (n = 7 mice, *t* = 2.839, *p* = 0.0236) and ACE-sdCKD mice (n = 11 mice, *t* = 7.104, *p* < 0.0001) exhibited cocaine-preferred behaviors. The ΔCPP score in ACE-dsC_KD_ mice is higher than that in ACE-sdC_Ctrl_ mice (n = 18 mice, *t* = 2.211, *p* = 0.0419), indicating an aggravating cocaine-preferred behavior in ACE-sdC_KD_ mice.

The percentage of c-Fos-positive & EGFP-positive in transfected CaMKII neurons was much higher in ACE-sdCKD mice than that in ACE-sdCCtrl mice (n = 16 slice from 8 mice, *t* = 4.744, *p* = 0.0003, Figure 4J). The ratio of p-ERK to ERK (n = 6 mice, *t* = 3.293, *p* = 0.0301), p-ERK (n = 6 mice, *t* = 3.903, *p* = 0.0175) levels were increased, and ERK (n = 6 mice, *t* = 0.1856, *p* = 0.8618) level was not changed by DUSP1 KD virus, compared to Ctrl virus (Figure 4K).

Together with the results of overexpressing DUSP1^CaMKII^, these results indicate that DUSP1^CaMKII^ mediate adolescent cocaine exposure-induced higher sensitivity to cocaine during adulthood, which is through regulating mPFC CaMKII-positive neuronal activation.

## Discussion

The current study found that ACE induce higher sensitivity to cocaine during adulthood. Subthreshold dose of cocaine (sdC), triggered more CaMKII-positive neurons in mPFC, especially subregions of Cg1 and PrL, accompanied with lower DUSP1 protein and higher *Dusp1* gene in mPFC of ACE mice during adulthood. To explore the roles of converse phenotypes of DUSP1 protein and *Dusp1* gene in response to sdC treatment, both upregulated and downregulated modulations of DUSP1^CaMKII^ were performed in mPFC of ACE mice. We found that specific overexpressing DUSP1^CaMKII^ efficiently blocked cocaine-induced CPP in ACE mice, and reduced CaMKII-positive neuronal activation in mPFC. By contrast, knocking-down DUSP1 aggravated cocaine-preferred behavior in ACE mice, and further activated CaMKII-positive neuronal activation in mPFC. ERK1/2 might be potential subsequent signal for DUSP1 in the process. (Figure 5).

**Figure 5.**
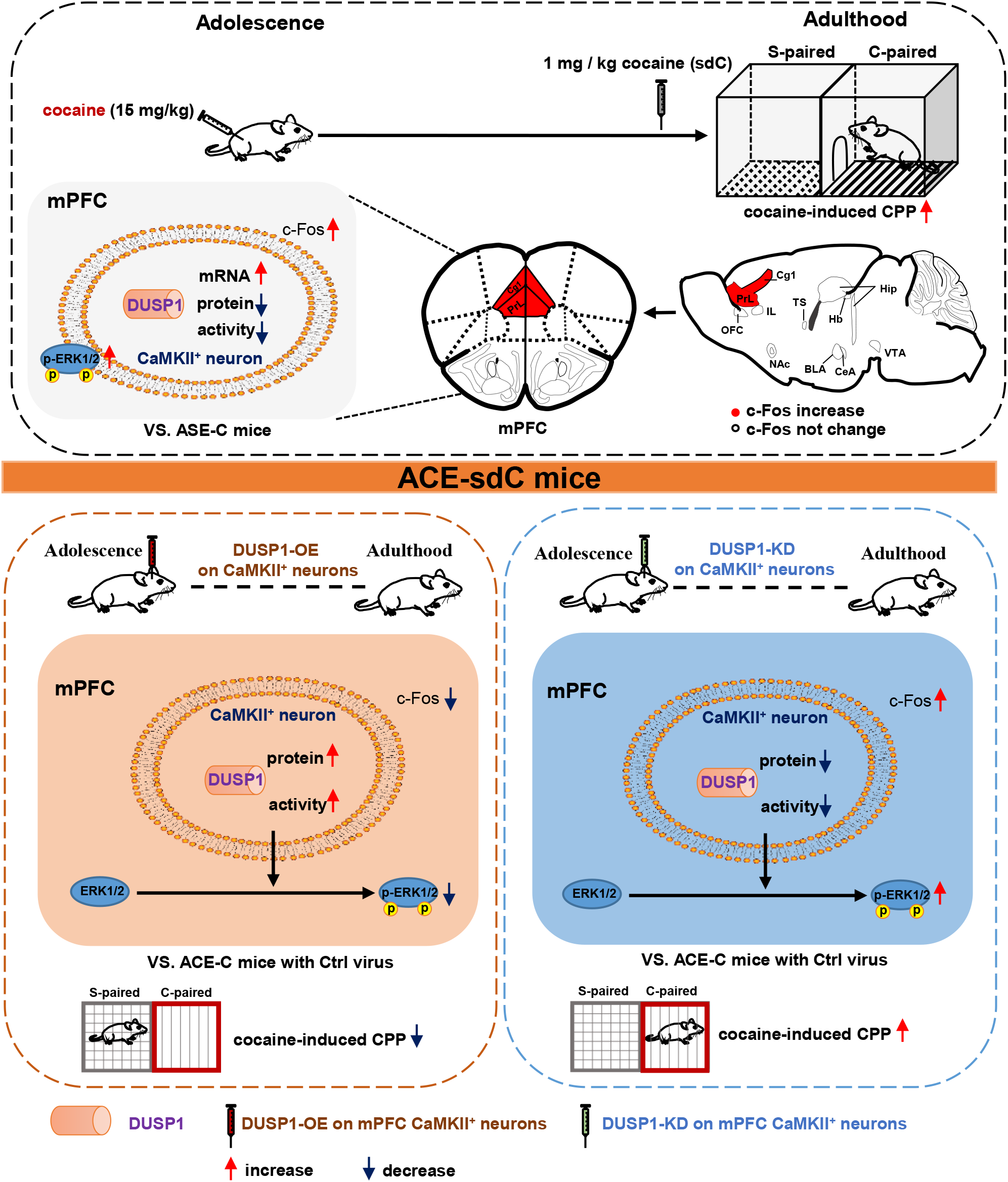
Schematic summary of the present study. Adolescent cocaine-exposed male mice (ACE) models were established by administrating cocaine of 15 mg / kg once daily during adolescent period. When growing to adult age, mice were subjected to conditioned place preference (CPP) to evaluate the sensitivity to cocaine. Subthreshold dose of cocaine (sdC), that is insufficient to produce CPP, was used to induce CPP in adulthood. The sdC treatment effectively induced CPP in ACE mice, but not in adolescent saline-exposed male mice (ASE) during adulthood, indicating higher sensitivity to cocaine caused by ACE. The sdC treatment triggered more CaMKII-positive neurons, and induced converse phenotypes of dual specificity phosphatase 1 (DUSP1) molecule in mPFC, as indicated by higher *Dusp1* gene, lower DUSP1 protein, lower DUSP1 activity and lower DUSP1 expression on CaMKII-positive neurons (DUSP1^CaMKII^) in mPFC of ACE mice than those of ASE mice during adulthood. To explore the role of converse phenotypes of DUSP1 protein and *Dusp1* gene in response to sdC treatment, both upregulated and downregulated modulations of DUSP1^CaMKII^ were performed in mPFC of ACE mice. Overexpressing DUSP1^CaMKII^ reduced CaMKII-positive neuronal activation, and ultimately blocked sdC-induced CPP in ACE mice during adulthood. While, knocking-down DUSP1^CaMKII^ activated more CaMKII-positive neurons, and aggravated sdC-preferred behavior in ACE mice during adulthood. ERK1/2 might be potential subsequent signal for DUSP1 in the process.

Adolescence is a critical period of brain development and maturation so that makes the adolescent brain highly susceptible to neurotoxicity of drugs. Exposure to Δ9-tetrahydrocannabinol (psychoactive ingredient of cannabis) enhanced sensitivity to cocaine in rats even into adulthood (24), and adolescent methylphenidate use (psychostimulant drug to treat attention deficit hyperactivity disorder) increased reactivity and vulnerability to cocaine in rats of adulthood (25), suggesting that adolescent use of the psychostimulant is an important risk factor for addiction susceptibility later in life. Previously using the similar ACE mice models with this study, we found that there was no difference on acquisition, extinction and reinstatement of CPP by repeated exposure to addictive doses of cocaine (15 mg/kg) in adulthood between ACE mice and controls (13). However, here we found that subthreshold dose administration of cocaine (1 mg/kg) effectively induced cocaine-preferred behaviors in ACE mice but not in control mice of their adulthood, indicating that ACE increase the sensitivity to cocaine which may render adult mice more susceptible to cocaine. Cass et.al (26) found that adolescent cocaine administration produced mPFC disinhibition that endured throughout adulthood. Our previous studies found that adolescent cocaine exposure induced prolonged modifications on synapses (14) and triggered neuronal activation (13) in mPFC of adult rodents, which contribute to the increased anxiety-like behaviors later in life. Here, by mapping the activation of potential brain regions involved in the increased sensitivity to cocaine, we found that mPFC, especially glutamatergic neurons in Cg1 and PrL, were obviously and strongly evoked in response to cocaine challenge in adult ACE mice. These findings suggest that ACE-induced plastic changes in mPFC play critical roles in facilitation of susceptibility to drugs later in life. Previously, we found that ACE suppressed mPFC activity partially through enhancing local GABAergic transmission on mPFC pyramidal neurons (13) and reduced dendritic spines of mPFC pyramidal neurons (14) in adulthood. However, unlike baseline condition, here we found that cocaine challenge in adulthood activated more glutamatergic neurons but did not change GABAergic including its subtypes of PV-positive and SOM-positive neurons in mPFC of ACE mice during adulthood. Based on these results, we hypothesized that the cocaine-changed signals in glutamatergic neurons of mPFC should be the mechanic basis for mPFC activation and susceptibility to cocaine-induced CPP during adulthood in mice treated with cocaine during adolescence.

In order to screen key signals, mPFC transcriptome was performed after cocaine administration in adult mice. We found that *Dusp1* gene and genes related to mitogen-activated protein kinase phosphatases (MAKPs) pathway were obviously upregulated in mPFC after cocaine challenge in ACE mice. While, level and activity of DUSP1 protein were decreased at the same time. We believe that the inconsistency between gene and protein changes of DUSP1 is due to their different meanings reflected in response to cocaine challenge. Takaki et al. (27) reported that *Dusp1* mRNA levels were significantly increased after acute or chronic methamphetamine administration in PFC of animals. We thought that the increased *Dusp1* gene in response to cocaine challenge, similar to immediate early genes, such as *c-fos, Jun, Ark*, reflected rapid and sensitive response of mPFC to cocaine in ACE mice, further indicating the important involvement of mPFC in process of susceptibility to drugs. In contrast, we found that the decreased DUSP1 protein level and activity was consistent with upregulated genes in MAPK-related signals by transcriptome analysis in mPFC of ACE mice by cocaine challenge during adulthood. Thus, DUSP1 protein might be the key regulatory molecule to plastic changes in mPFC which underlies the increased sensitivity to drugs in ACE mice during adulthood. Indeed, we found that specific overexpression but not knockdown of mPFC DUSP1^CaMKII^ efficiently suppressed cocaine-activated mPFC, and ultimately reduced cocaine-preferred behaviors in ACE mice. As a negative regulatory factor on MAPK-related signals, DUSP1 is also known as MKP-1. By dephosphorylating p38, JNKs and ERK (28), DUSP1 regulates the brain function. Among them, ERK-related signals are well-studied in cocaine addiction. The ERK signaling pathway is a canonical MAPK signaling pathway that has been implicated in synaptic plasticity and neuronal growth (29). Acute cocaine administration is sufficient to increased phosphorylation of ERK in PFC, which contribute to the long-term plastic changes by cocaine in the brain (30–32). While, blockade of ERK activation during re-exposure to cocaine erased previously cocaine-preferred behaviors (33). In the present study, we found that levels of p-ERK increased when some changes in mPFC by cocaine, while it was decreased in DUSP1-overexpressed mPFC with reduced activity in ACE mice, suggesting that DUSP1 may recruit ERK-related signals to regulate mPFC activity, and therefore influence sensitivity to cocaine challenge in ACE mice. In the future study, the mechanistic investigation of DUSP1/MAPK signal pathway underlying the phenotypes of higher cocaine-preferred behaviors should be further explored in ACE mice.

Collectively, our findings reveal a novel role of mPFC DUSP1^CaMKII^ in higher sensitivity to cocaine in ACE mice during adulthood, which is through regulating mPFC activity. The mPFC DUSP1^CaMKII^ may represent a promising therapeutic target for treatment of addiction, especially caused by adolescent substance use.

## Materials and methods

### Animals

Male C57BL/6 wild type (WT) mice weighing 12-15 g were used. All animals were housed at constant humidity (40~60%) and temperature (24 ± 2°C) with a 12-hour light/dark cycle (lights on at 8 a.m.) and allowed free access to food and water. All mice were handled for three days before onset of experiments. All procedures were carried out in accordance with the National Institutes of Health Guide for the Care and Use of Laboratory Animals and approved by the Institutional Animal Care and Use Committee (IACUC) at Nanjing University of Chinese Medicine.

### Drug treatment

From P28 to P42, male C57BL/6 WT mice were assigned to receive cocaine hydrochloride (15 mg/kg, dissolved in saline, i.p., Qinghai Pharmaceutical, China) or saline (0.2 mL, i.p.) once daily for 15 consecutive days. From P43-P74, these mice were kept at their home cage (4 mice per cage). From P75 to P78, a subset of mice were subjected to CPP training and test induced by subthreshold dose of cocaine hydrochloride (1 mg/kg).

### CPP

The CPP test was performed in the TopScan3D CPP apparatus (CleverSys, VA, USA), which is constructed of two distinct chambers (15 × 15 × 23 cm each) separated by a removable guillotine door. The CPP procedure consisted of three phases: the pre-conditioning test (Pre-test, P75), conditioning (CPP training, P76-77) and post-conditioning test (Test, P78). Here, a subthreshold dose (1 mg/kg) of cocaine, which is believed not enough to induce CPP in naïve mouse, was used to induce CPP in mice on P78. During CPP training, mice were injected with saline (0.2 mL, i.p.) in the morning and cocaine (1 mg/kg, i.p.) in the afternoon once daily. After each injection, the mice were confined to one chamber (non-drug-paired chamber or drug-paired chamber) for 45 min. During the Pre-test and Test, mice freely access two chambers for 15 min. The CPP score is the time spent in drug-paired chamber minus that in non-drug-paired chamber, and the ΔCPP score is the test CPP score minus the pre-test CPP score.

### Immunofluorescence

On P78, brain tissue were collected from a subset of mice within 90 min after CPP test. Some brains of those were perfused with 4% paraformaldehyde (PFA). The coronal brain sections (30 μm) were cut on a cryostat (Leica, Germany). The sections were incubated in 0.3% (v/v) Triton X-100 for 0.5 h, blocked with 5% donkey serum for 1.5 h at room temperature, and incubated overnight at 4°C with the following primary antibodies: rabbit anti-c-Fos (1:1500, RRID: AB_2247211, Cell Signalling Technology, USA), mouse anti-NeuN (1:800, RRID: AB_2298772, Millipore, USA), DUSP1 (1:100, BM4856, Boster Biotechnology, China), mouse anti-GAD67 (1:500, RRID: AB_2278725, Millipore, USA), mouse anti-PV (1:250, BM1339, Boster Biotechnology, China), mouse anti-SOM (1:50, RRID:AB_2271061, Santa Cruz, USA), mouse anti-CaMKIIα (1:100, SC-13141, Santa Cruz, USA) and mouse anti-CaMKIIα (1:200, RRID: AB_2721906, Cell Signalling Technology, USA), followed by the corresponding fluorophore-conjugated secondary antibodies for 1.5 h at room temperature. The following secondary antibodies were used here: Alexa Fluor 555-labeled donkey anti-rabbit secondary antibody (1:500, RRID: AB_2762834, Invitrogen, USA), Alexa Fluor 488-labeled donkey anti-mouse secondary antibody (1:500, RRID: AB_141607, Invitrogen, USA). Fluorescence signals were visualized using a Leica DM6B upright digital research microscope (Leica, Germany) or Leica TCS SP8 (Leica, Germany).

### Fiber photometry

On P51, the *rAAV2/9-CaMKII-GCaMp6m* (PT-0111, 3.24E+12 vg/ml, Brain VTA, China) virus was bilaterally injected into mPFC (AP + 2.10 mm, ML ± 0.3 mm, DV – 2.15 mm), especially into PrL of mice. On P78, the GCaMp6m signals in mPFC were recorded during CPP test. An optical fiber (200 μm outer diameter, 0.37 numerical aperture, Inper Ltd., China) was placed 100 μm above the viral injection site. The calcium-dependent fluorescence signals were obtained by stimulating cells that transfected GCaMp6m virus with a 470 nm LED (35-40 μW at fiber tip), while calcium-independent signals were obtained by stimulating these cells with a 405 nm LED (15-20 μW at fiber tip). The LED lights of 470 nm and 405 nm were alternated at 66 fps and light emission was recorded using sCMOS camera containing the entire fiber bundle (2 m in length, NA = 0.37, 200 μm core, Inper Ltd.). The analog voltage signals fluorescent was filtered at 30 Hz and digitalized at 100 Hz. The GCaMp6m signals were recorded and analyzed by Inper Studio Multi Color EVAL15 software (Inper Ltd., China) and Inper Data Process V0.5.9 (Inper Ltd., China), respectively. F0 is the baseline fluorescence signal which was recorded for 1 sec before mice entering into the cocaine-paired chamber. F is the real-time fluorescence signal which was recorded for 4 sec after mice entering into the cocaine-paired chamber during CPP test. Each recording trail was with 5 sec recording duration. The values of ΔF/F are calculated by (F-F0)/F0. The area under curve (AUC) is the integral under recording duration related to corresponding baseline at every trial.

### The mRNA sequencing and data processing in mPFC

On P78, mPFC tissue (n = 6 per group) were collected from mice after CPP test. Library preparation was performed by Novogene Bioinformatics Technology Co., Ltd (China). RNA degradation and purity were monitored on 1% agarose gels and using the NanoPhotometer ^®^ spectrophotometer (IMPLEN, CA, USA). RNA integrity was assessed using the RNA Nano 6000 Assay Kit of the Bioanalyzer 2100 system (Agilent Technologies, CA, USA). The mRNA was purified and fragmented from a total amount of 1 μg RNA per sample. The cDNA was synthesized, adenylated and ligated with hairpin-structured NEBNext Adaptor before hybridization. 250-300 bp cDNA fragments were purified and preferentially selected for hybridization. The PCR products were purified (AMPure XP system) and library quality was assessed on the Agilent Bioanalyzer 2100 system. Index-coded samples were clustered on a cBot Cluster Generation System using TruSeq PE Cluster Kit v3-cBot-HS (Illumia) and sequenced on an Illumina Novaseq platform. Differential mRNA expression analysis was performed using the DESeq2 R package (1.16.1). The *p* values were adjusted using the Benjamini & Hochberg method. Cutoffs of an absolute fold change > 1 and a correlated *p* value < 0.05 were set for significantly differential expression. The Bioinformatics analysis, including GO, KEGG and PPI of upregulated genes by a subthreshold dose of cocaine in ACE mice were performed in this study.

### RNA extraction and quantitative real-time PCR

On P78, brain tissue were collected from a subset of mice within 90 min after CPP test. The mPFC tissue were dissected immediately on dry ice. Total RNA was extracted from mPFC using FastPure Cell/Tissue Total RNA Isolation Kit (Vazyme, China). After RNA extraction, the amount of RNA was normalized across samples, and cDNA was created using Vazyme (HiScript II Q RT SuperMix for Qpcr (+gDNA wiper)). The β-actin was used as the internal control. The primers of DUSP1 and β-actin were as follows: DUSP1 (forward, GTTGTTGGATTGTCGCTCCTT; reverse, TTGGGCACGATATGCTCCAG), β-actin (forward, GGCTGTATTCCCCTCCATCG; reverse, CCAGTTGGTAACAATGCCATGT). DUSP1 mRNA levels were normalized to β-actin mRNA levels. The relative mRNA level was calculated by the comparative CT method (2^-ΔΔCt^).

### Western blot

On P74 and P78, brain tissue were collected from two subsets of mice without CPP training (P74) or within 90 min after CPP test (P78). The mPFC tissue were dissected immediately on dry ice. Total mPFC protein was extracted using RIPA lysis buffer (Shanghai Beyotime, China). Protein samples (15 μg) was separated by 10% SDS–PAGE or 12% SDS–PAGE and electrophoretically transferred onto PVDF membranes. The transferred membranes were blocked with 5% non-fat dry milk and 0.1% Tween 20 in 10 mM Tris–HCl (TBST buffer, pH 7.5) for 1.5 h at room temperature, then subsequently incubated with the following primary antibodies: c-Fos (1:1500, Rabbit, RRID: AB_2247211, Cell Signaling Technology, USA), DUSP1 (1:1000, Rabbit, BM4856, Boster Biotechnology, China), ERK_1/2_ (1:1000, Rabbit, BM4326, Boster Biotechnology, China), p-ERK_1/2_ (1:1000, Rabbit, BM4156, Boster Biotechnology, China). The next day, the membranes were washed three times in Tris-buffered saline with Tween 20 (TBS-T) and incubated with horseradish peroxidase (HRP)-conjugated secondary antibody goat anti-rabbit (1:5000, RRID: AB_2814709, Beijing ComWin Biotech Co., China) at room temperature for 1 h. The blots were visualized by the ECL kit (Beijing ComWin Biotech Co., China) and the signal was visualized by imaging system (Tanon-5200, Shanghai, China). The blots were washed with stripping buffer (Beyotime Institute of Biotechnology) to allow reprobing with other antibodies. In this study, β-actin was used as control. Values for target protein levels were calculated using Image J software (NIH, USA). The relative level of each protein was normalized to β-actin.

### Total dual specificity phosphatase 1 (DUSP1) measurement in mPFC by ELISA

On P74 and P78, brain tissue were collected from two subsets of mice without CPP training (P74) or within 90 min after CPP test (P78). The mPFC tissue were dissected immediately on dry ice. The activity of mPFC DUSP1 was measured by commercial ELISA (Mouse DUSP1 ELISA Kit instruction, Shanghai shyx-bio.com, China). The mPFC tissues were diluted 1:10 in saline and mashed using Tissue Grinder (Wuhan Servicebio technology CO., China), then centrifuged for 30 min at 3000×g at 2-8 °C and the supernatants were store samples at −20 °C. The 50 μL mixture of sample and Sample Diluent (1:4) were put in one well of plate. Then, add 100 μL of HRP-conjugate reagent to each well, cover with an adhesive strip and incubate for 60 min at 37°C. Washing the wells with Wash Solution (400 μL) five times. Then, added chromogen solution A 50 μL and chromogen solution B 50 μL to each well and incubated for 15 min at 37°C. The reaction was stopped by adding 50 μL Stop Solution. Read the Optical Density (O.D.) at 450 nm using a standard microplate reader (BIO-RAD iMark Microplate Reader, USA) within 15 min.

### Specific overexpression or knockdown of mPFC DUSP1^CaMKII^

On P51, four cohorts of mice were anesthetized with 2% isoflurane in oxygen, and were fixed in a stereotactic frame (RWD, China). A heating pad was used to maintain the core body temperature of the animals at 36°C. The coordinates of mPFC were AP + 2.10, ML ± 0.3, DV – 2.15 in mm. In Experiment of overexpression of DUSP1, a volume of 400 nL of *CaMKIIap-MCS-EGFP-3Flag-SV40 PloyA (AAV9-CaMKIIap-DUSP1-RNAOE-EGFP*, 4.90E+13v.g/ml; Cat#: GOSV0322969, Gene Chem, China) or *CaMKIIap-EGFP-SV40 PloyA (AAV9-CaMKIIap-EGFP*, 4.01E+12v.g/ml; Cat#: AAV9/CON540, Gene Chem, China) bilaterally into the mPFC (AP + 2.10 mm, ML ± 0.3 mm, DV – 2.15 mm) at a rate of 80 nL/min. In Experiment of knockdown of DUSP1, a volume of 400 nL of *CaMKIIap-EGFP-MIR155(MCS)-SV40 PloyA (AAV9-CaMKIIap-DUSP1-RNAi-EGFP*, 1.10E+13 v.g/ml; Cat#: GIDV0295686, Gene Chem, China) or *CaMKIIap-EGFP-SV40 PloyA (AAV9-CaMKIIap-EGFP*, 4.01E+12v.g/ml; Cat#: AAV9/CON540, Gene Chem, China) bilaterally into the at a rate of 80 nL/min. After surgery, mice were maintained at home cage about 3 weeks.

### Statistical analysis

Statistical analysis was carried out using GraphPad Prism 8.0.2 software. All data are presented as the Mean ± SEM. The data of CPP score between ASE-C and ACE-C groups are analyzed by two-way analysis of variance (ANOVA), while CPP score between Pre-test and Test in each mouse were analyzed by paired *t*-tests. Other data were analyzed by unpaired *t*-tests. GO and KEGG enrichment analysis of differentially expressed genes was implemented by the clusterProfiler R package, and a corrected *p* < 0.05 were considered as significantly enriched. The PPI network was drawn via the cytoscape 3.4.0 analysis software. All statistical significance was set as *p* < 0.05.

## Author contributions

Wei X and Chang J have equal contribution to the manuscript. Wei X, Chang J, Cheng Z and Fan Y performed behavioral tests, morphological tests and virus-related experiment. Chen W, Guo H, Liu Z, Mai Y, Hu T, Zhang Y and Cai Q assist the data analysis and organized figures. Fan Y, Chang J and Ge F performed data analysis. Guan X and Fan Y wrote the manuscript. Guan X developed the overall concept.

## Acknowledgments

This work is supported by National Natural Science Foundation of China (82271531 and 82071495), Natural Science Foundation of Jiangsu Province, China (BK20201398) and Natural Science Foundation of the Higher Education Institutions of Jiangsu Province, China (21KJB360007).

## Data availability statement

The data that support the findings of this study are available from the corresponding author upon reasonable request.

## References

1. Chambers RA, et al. Developmental neurocircuitry of motivation in adolescence: a critical period of addiction vulnerability. The American journal of psychiatry. 2003;160(6):1041–52.

2. Spear LP. Consequences of adolescent use of alcohol and other drugs: Studies using rodent models. Neuroscience and biobehavioral reviews. 2016;70(228–43.

3. Wong WC, and Marinelli M. Adolescent-onset of cocaine use is associated with heightened stress-induced reinstatement of cocaine seeking. Addiction biology. 2016;21(3):634–45.

4. Jordan CJ, and Andersen SL. Sensitive periods of substance abuse: Early risk for the transition to dependence. Developmental Cognitive Neuroscience. 2017;25(29–44.

5. Perry JL, et al. Prefrontal cortex and drug abuse vulnerability: translation to prevention and treatment interventions. Brain Res Rev. 2011;65(2):124–49.

6. Zhang Y, et al. Molecular changes in the medial prefrontal cortex and nucleus accumbens are associated with blocking the behavioral sensitization to cocaine. Scientific reports. 2015;5(1):16172.

7. Koss WA, et al. Dendritic remodeling in the adolescent medial prefrontal cortex and the basolateral amygdala of male and female rats. Synapse. 2014;68(2):61–72.

8. Shapiro LP, et al. Differential expression of cytoskeletal regulatory factors in the adolescent prefrontal cortex: Implications for cortical development. J Neurosci Res. 2017;95(5):1123–43.

9. Caffino L, et al. A single cocaine administration alters dendritic spine morphology and impairs glutamate receptor synaptic retention in the medial prefrontal cortex of adolescent rats. Neuropharmacology. 2018;140(209–16.

10. Caffino L, et al. A single cocaine exposure disrupts actin dynamics in the cortico-accumbal pathway of adolescent rats: modulation by a second cocaine injection. Psychopharmacology. 2017;234(8):1217–22.

11. Chocyk A, et al. Early-life stress affects the structural and functional plasticity of the medial prefrontal cortex in adolescent rats. The European journal of neuroscience. 2013;38(1):2089–107.

12. Fuhrmann D, et al. Adolescence as a Sensitive Period of Brain Development. Trends in cognitive sciences. 2015;19(10):558–66.

13. Shi P, et al. Adolescent cocaine exposure enhances the GABAergic transmission in the prelimbic cortex of adult mice. FASEB journal: official publication of the Federation of American Societies for Experimental Biology. 2019;33(7):8614–22.

14. Zhu W, et al. Adolescent cocaine exposure induces prolonged synaptic modifications in medial prefrontal cortex of adult rats. Brain structure & function. 2018;223(4):1829–38.

15. Low HB, and Zhang Y. Regulatory Roles of MAPK Phosphatases in Cancer. Immune network. 2016;16(2):85–98.

16. Wang X, et al. DUSP1 Promotes Microglial Polarization toward M2 Phenotype in the Medial Prefrontal Cortex of Neuropathic Pain Rats via Inhibition of MAPK Pathway. ACS Chemical Neuroscience. 2021;12(6):966–78.

17. Xia L, et al. Rosiglitazone Improves Glucocorticoid Resistance in a Sudden Sensorineural Hearing Loss by Promoting MAP Kinase Phosphatase-1 Expression. Mediators of inflammation. 2019;2019(7915730.

18. Taylor DM, et al. MAP kinase phosphatase 1 (MKP-1/DUSP1) is neuroprotective in Huntington’s disease via additive effects of JNK and p38 inhibition. The Journal of neuroscience: the official journal of the Society for Neuroscience. 2013;33(6):2313–25.

19. Xu P, et al. DUSP1 alleviates cerebral ischaemia reperfusion injury via inactivating JNK-Mff pathways and repressing mitochondrial fission. Life Sci. 2018;210(251–62.

20. Ndong C, et al. Mitogen activated protein kinase phosphatase-1 prevents the development of tactile sensitivity in a rodent model of neuropathic pain. Molecular pain. 2012;8(34.

21. Jefsen OH, et al. Transcriptional regulation in the rat prefrontal cortex and hippocampus after a single administration of psilocybin. Journal of Psychopharmacology. 2020;35(4):483–93.

22. Wojcieszak J, et al. Induction of immediate early genes expression in the mouse striatum following acute administration of synthetic cathinones. Pharmacological Reports. 2019;71(6):977–82.

23. Spear LP. The adolescent brain and age-related behavioral manifestations. Neuroscience and biobehavioral reviews. 2000;24(4):417–63.

24. Friedman AL, et al. Effects of adolescent Δ9-tetrahydrocannabinol exposure on the behavioral effects of cocaine in adult Sprague-Dawley rats. Experimental and clinical psychopharmacology. 2019;27(4):326–37.

25. Brandon CL, et al. Enhanced reactivity and vulnerability to cocaine following methylphenidate treatment in adolescent rats. Neuropsychopharmacology: official publication of the American College of Neuropsychopharmacology. 2001;25(5):651–61.

26. Cass DK, et al. Developmental disruption of gamma-aminobutyric acid function in the medial prefrontal cortex by noncontingent cocaine exposure during early adolescence. Biological psychiatry. 2013;74(7):490–501.

27. Takaki M, et al. Two kinds of mitogen-activated protein kinase phosphatases, MKP-1 and MKP-3, are differentially activated by acute and chronic methamphetamine treatment in the rat brain. Journal of neurochemistry. 2001;79(3):679–88.

28. Lawan A, et al. Skeletal Muscle–Specific Deletion of MKP-1 Reveals a p38 MAPK/JNK/Akt Signaling Node That Regulates Obesity-Induced Insulin Resistance. Diabetes. 2018;67(4):624–35.

29. Sweatt JD. The neuronal MAP kinase cascade: a biochemical signal integration system subserving synaptic plasticity and memory. Journal of neurochemistry. 2001;76(1):1–10.

30. Valjent E, et al. Addictive and non-addictive drugs induce distinct and specific patterns of ERK activation in mouse brain. The European journal of neuroscience. 2004;19(7):1826–36.

31. Radwanska K, et al. Regulation of cocaine-induced activator protein 1 transcription factors by the extracellular signal-regulated kinase pathway. Neuroscience. 2006;137(1):253–64.

32. Sun WL, et al. Molecular Mechanism: ERK Signaling, Drug Addiction, and Behavioral Effects. Progress in molecular biology and translational science. 2016;137(1–40.

33. Valjent E, et al. Inhibition of ERK pathway or protein synthesis during reexposure to drugs of abuse erases previously learned place preference. Proceedings of the National Academy of Sciences of the United States of America. 2006;103(8):2932–7.

