## Supplementary material for "The DUSP1 on CaMKII-positive neurons in mPFC mediates adolescent cocaine exposure-induced higher sensitivity to cocaine during adulthood": Suppl legends and figures

**Supplementary Data**

**Supplementary Figure legends**

**Figure S1. The number of c-Fos-positive neurons in brain areas. A,** The number of c-Fos-positive neurons in brain areas of ASE-sdC and ACE-sdC mice. OFC, orbitofrontal cortex; Cg1, cingulate cortex; PrL, prelimbic cortex; IL, infralimbic cortex; NAc, nucleus accumbens; BNST, bed nucleus of the stria terminalis; TS, triangular septal nucleus; BLA, basolateral amygdala; CeA, central nucleus of the amygdala; mHb, medial habenula; LHb, lateral habenular; dCA1, vCA1, dorsal and ventral hippocampal CA1; dCA3, dorsal hippocampal CA3; dDG, dorsal dentate gyrus; VTA, ventral tegmental area. ASE-sdC, adolescent saline-exposed mice with subthreshold dose of cocaine treatment during adulthood; ACE-sdC, adolescent cocaine-exposed mice with subthreshold dose of cocaine treatment during adulthood; N.S., $p > 0.05$ vs ASE-sdC mice.

**Figure S2. The c-Fos levels and staining in sublayers of mPFC subregions. A,** The protein levels of c-Fos on P74. **B,** The number of c-Fos-positive CaMKII-labeled, GAD67-labeled, PV-labeled or SOM-labeled neurons in mPFC. ASE-sdC, adolescent saline-exposed mice with subthreshold dose of cocaine treatment during adulthood; ACE-sdC, adolescent cocaine-exposed mice with subthreshold dose of cocaine treatment during adulthood; N.S., $p > 0.05$, **, $p < 0.01$ vs ASE-sdC mice.

**Figure S3. The mPFC gene expression in adult mice. A,** Volcano plot illustration showed the upregulated, downregulated and unchanged genes in response to cocaine administration in adult mice. Each dot represents one gene. Red, blue and gray dots represent the significantly up-regulated, down-regulated and not changed genes. **B,** The Protein-Protein Interaction (PPI) network of up-regulated genes. **C,** The heatmaps of up-regulated genes. The gene clusters, which are initially the sample type, are presented along the horizontal axis. The threshold was set as *p*-value < 0.05, single sample gene expression amount exceeds 100. Columns represent samples, and rows show the different genes. Color scale, red for higher abundance, while purple for lower abundance. **D,** Top 30 of enriched GO terms of upregulated genes by the categories of BP, CC and MF. **E,** Top 20 of enriched KEGG pathways of upregulated genes. ASE-sdC, adolescent saline-exposed mice with subthreshold dose of cocaine treatment during adulthood; ACE-sdC, adolescent cocaine-exposed mice with subthreshold dose of cocaine treatment during adulthood.

### Figure S1

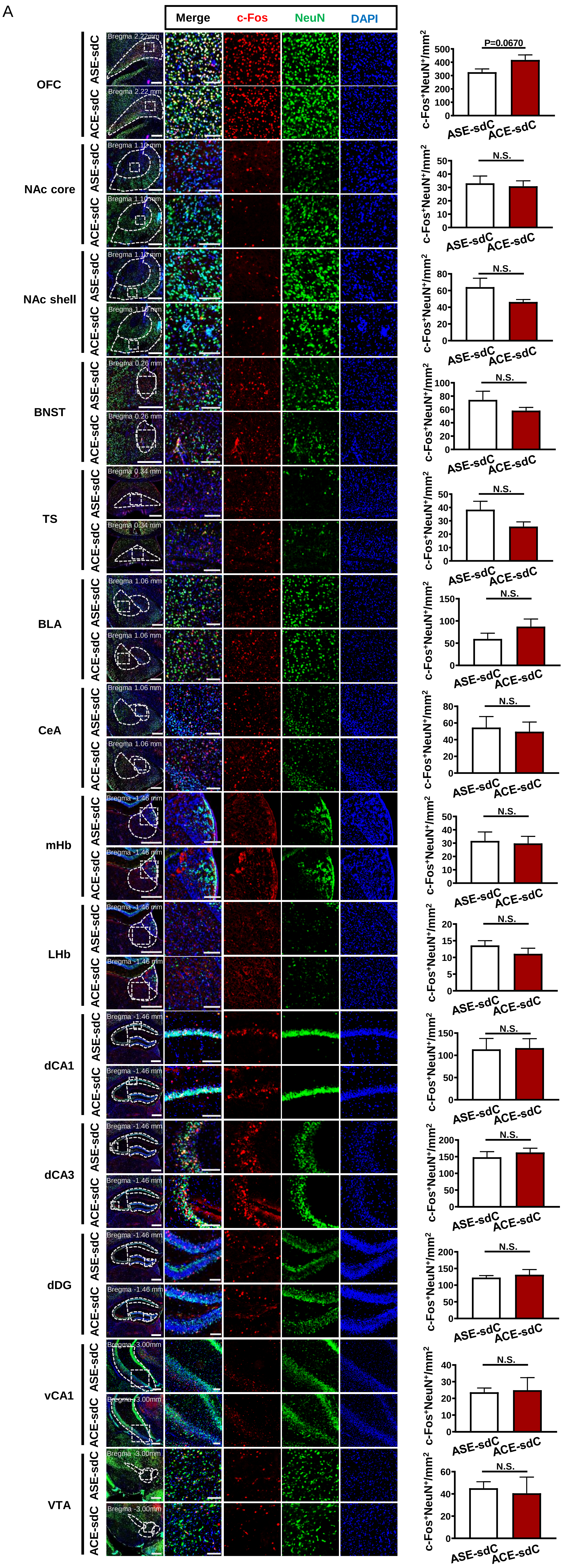
A
Merge
c-Fos
NeuN
DAPI
OFC
NAc core
NAc shell
BNST
TS
BLA
CeA
mHb
LHb
dCA1
dCA3
dDG
vCA1
VTA
ASE-sdC
ACE-sdC
Bregma 2.22 mm
Bregma 1.10 mm
Bregma 0.26 mm
Bregma 0.34 mm
Bregma 1.06 mm
Bregma -1.46 mm
Bregma -3.00mm
c-Fos+NeuN+/mm2
P=0.0670
N.S.

**Figure S2**

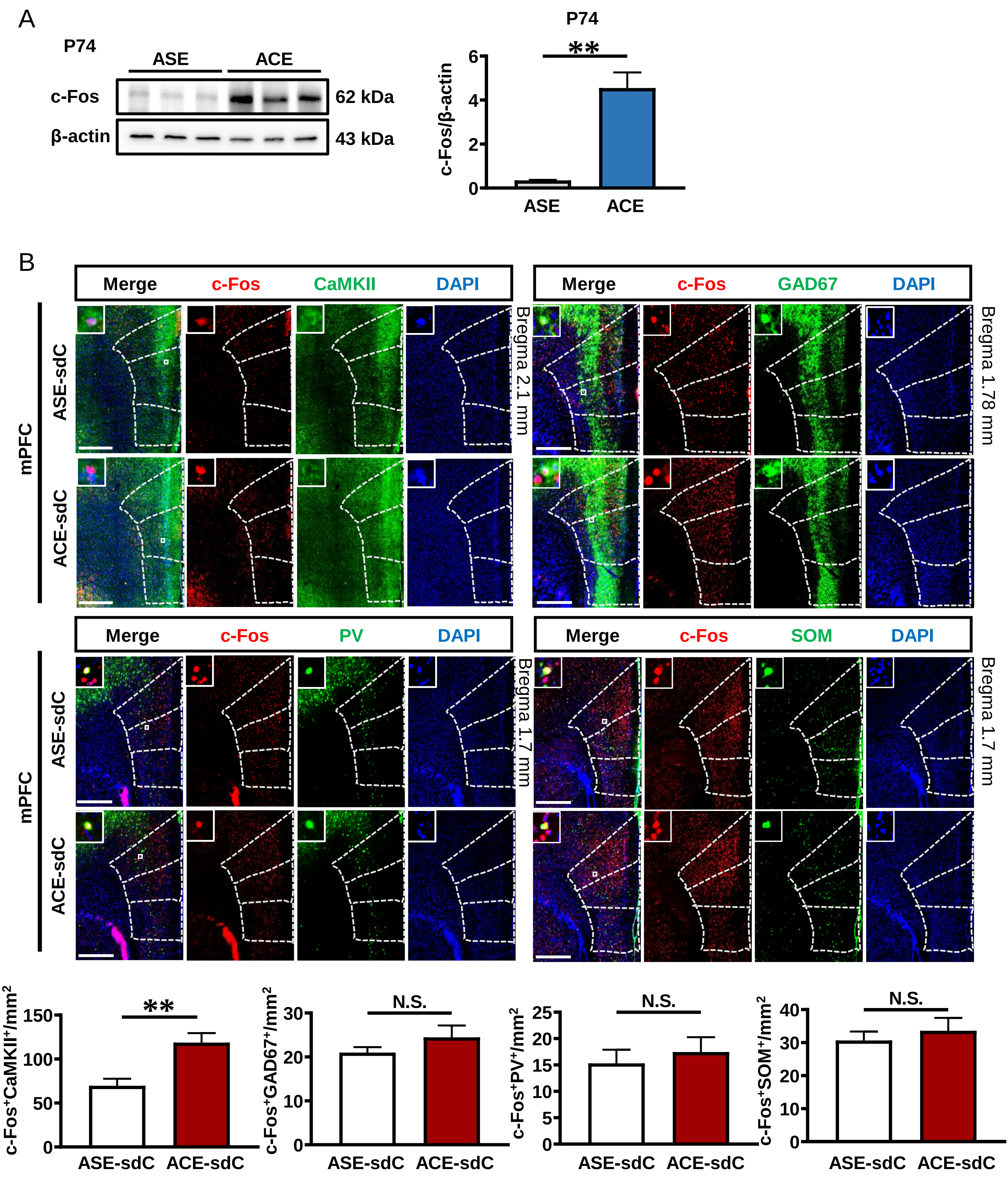
A
P74
ASE
ACE
c-Fos
β-actin
62 kDa
43 kDa
P74
**
c-Fos/β-actin
ASE
ACE
B
Merge
c-Fos
CaMKII
DAPI
GAD67
PV
SOM
mPFC
ASE-sdC
ACE-sdC
Bregma 2.1 mm
Bregma 1.78 mm
Bregma 1.7 mm
c-Fos+CaMKII+/mm2
c-Fos+GAD67+/mm2
c-Fos+PV+/mm2
c-Fos+SOM+/mm2
N.S.

Figure S3

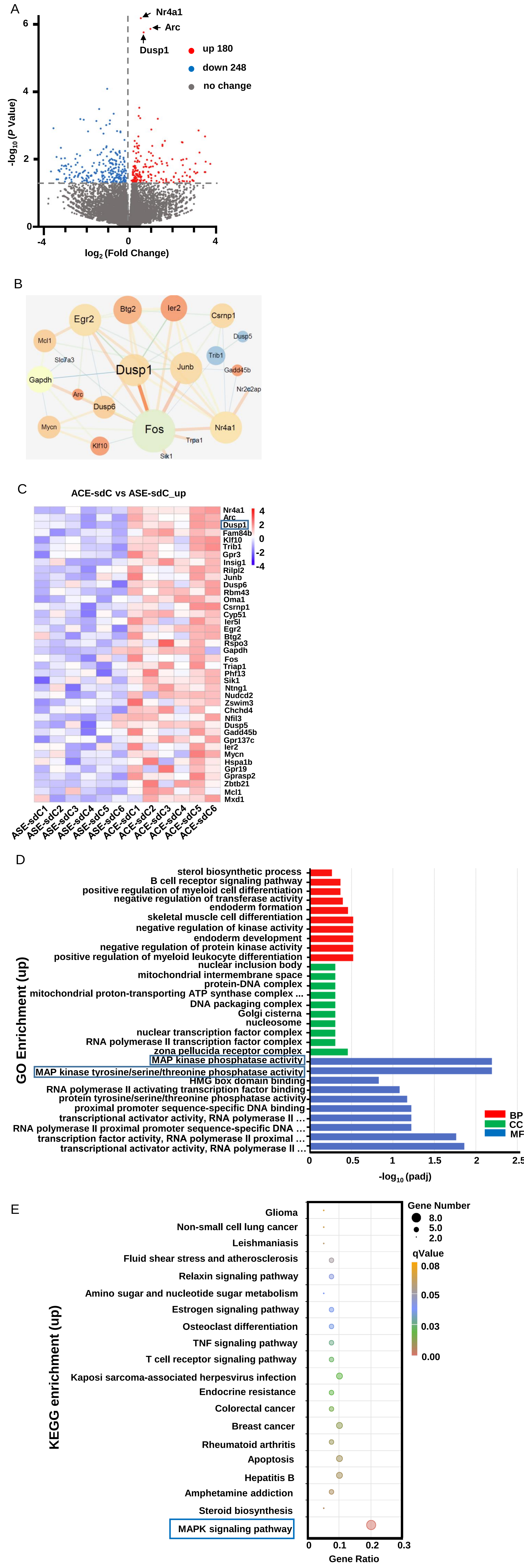

A
Nr4a1
Arc
Dusp1
up 180
down 248
no change
-log10 (P Value)
log2 (Fold Change)
0
2
4
6
-4
0
4
B
Btg2
Ier2
Egr2
Csrnp1
Mcl1
Dusp5
Slc7a3
Trib1
Gapdh
Dusp1
Junb
Gadd45b
Nr2c2ap
Arc
Dusp6
Mycn
Fos
Nr4a1
Klf10
Trpa1
Sik1
C
ACE-sdC vs ASE-sdC_up
Nr4a1
Arc
Dusp1
Fam84b
Klf10
Trib1
Gpr3
Insig1
Rilpl2
Junb
Dusp6
Rbm43
Oma1
Csrnp1
Cyp51
Ier5l
Egr2
Btg2
Rspo3
Gapdh
Fos
Triap1
Phf13
Sik1
Ntng1
Nudcd2
Zswim3
Chchd4
Nfil3
Dusp5
Gadd45b
Gpr137c
Ier2
Mycn
Hspa1b
Gpr19
Gprasp2
Zbtb21
Mcl1
Mxd1
4
2
0
-2
-4
ASE-sdC1
ASE-sdC2
ASE-sdC3
ASE-sdC4
ASE-sdC5
ASE-sdC6
ACE-sdC1
ACE-sdC2
ACE-sdC3
ACE-sdC4
ACE-sdC5
ACE-sdC6
D
sterol biosynthetic process
B cell receptor signaling pathway
positive regulation of myeloid cell differentiation
negative regulation of transferase activity
endoderm formation
skeletal muscle cell differentiation
negative regulation of kinase activity
endoderm development
negative regulation of protein kinase activity
positive regulation of myeloid leukocyte differentiation
nuclear inclusion body
mitochondrial intermembrane space
protein-DNA complex
mitochondrial proton-transporting ATP synthase complex ...
DNA packaging complex
Golgi cisterna
nucleosome
nuclear transcription factor complex
RNA polymerase II transcription factor complex
zona pellucida receptor complex
MAP kinase phosphatase activity
MAP kinase tyrosine/serine/threonine phosphatase activity
HMG box domain binding
RNA polymerase II activating transcription factor binding
protein tyrosine/serine/threonine phosphatase activity
proximal promoter sequence-specific DNA binding
transcriptional activator activity, RNA polymerase II ...
RNA polymerase II proximal promoter sequence-specific DNA ...
transcription factor activity, RNA polymerase II proximal ...
transcriptional activator activity, RNA polymerase II ...
GO Enrichment (up)
0
0.5
1
1.5
2
2.5
-log10 (padj)
BP
CC
MF
E
Glioma
Non-small cell lung cancer
Leishmaniasis
Fluid shear stress and atherosclerosis
Relaxin signaling pathway
Amino sugar and nucleotide sugar metabolism
Estrogen signaling pathway
Osteoclast differentiation
TNF signaling pathway
T cell receptor signaling pathway
Kaposi sarcoma-associated herpesvirus infection
Endocrine resistance
Colorectal cancer
Breast cancer
Rheumatoid arthritis
Apoptosis
Hepatitis B
Amphetamine addiction
Steroid biosynthesis
MAPK signaling pathway
KEGG enrichment (up)
0
0.1
0.2
0.3
Gene Ratio
Gene Number
8.0
5.0
2.0
qValue
0.08
0.05
0.03
0.00
